# Social Selection and the Evolution of Maladaptation

**DOI:** 10.1101/2021.03.12.435141

**Authors:** Joel W. McGlothlin, David N. Fisher

## Abstract

Evolution by natural selection is often viewed as a process that inevitably leads to adaptation, or an increase in population fitness over time. However, maladaptation, an evolved decrease in fitness, may also occur in response to natural selection under some conditions. Social selection, which arises from the effects of social partners on fitness, has been identified as a potential cause of maladaptation, but we lack a general rule identifying when social selection should lead to a decrease in population mean fitness. Here we use a quantitative genetic model to develop such a rule. We show that maladaptation is most likely to occur when social selection is strong relative to nonsocial selection and acts in an opposing direction. In this scenario, the evolution of traits that impose fitness costs on others may outweigh evolved gains in fitness for the individual, leading to a net decrease in population mean fitness. Further, we find that maladaptation may also sometimes occur when phenotypes of interacting individuals negatively covary. We outline the biological situations where maladaptation in response to social selection can be expected, provide both quantitative genetic and phenotypic versions of our derived result, and suggest what empirical work would be needed to test it. We also consider the effect of social selection on inclusive fitness and support previous work showing that inclusive fitness cannot suffer an evolutionary decrease. Taken together, our results show that social selection may decrease population mean fitness when it opposes individual-level selection, even as inclusive fitness increases.

**Subject area:** Quantitative genetics and Mendelian inheritance

## Introduction

Adaptive evolution is often visualized as a hill-climbing process, with natural selection causing populations to ascend peaks of higher fitness on an adaptive landscape (Wright 1932; Simpson 1944; Lande 1976, 1979; Arnold et al. 2001; Estes and Arnold 2007; Svensson and Calsbeek 2012; Hendry 2017). The adaptive landscape view implies that in the absence of constraints, natural populations should tend to be either well adapted to their current environment, i.e., occupying a local fitness peak, or adapting, i.e., climbing a local fitness peak. Directional selection has been shown to be particularly common in nature, suggesting that populations are indeed often adapting to stable or changing environmental conditions (Hoekstra et al. 2001; Kingsolver et al. 2001; Hereford et al. 2004; Siepielski et al. 2009, 2013; Kingsolver and Diamond 2011; Morrissey and Hadfield 2012; Hendry 2017). More limited evidence for stabilizing selection suggests that natural populations are sometimes at or near fitness peaks, at least for some traits (Kingsolver et al. 2001; Estes and Arnold 2007; Kingsolver and Diamond 2011). Under certain circumstances, however, populations may persist in a state of suboptimal mean fitness or may even be undergoing a process of evolved reduction in mean fitness. The term “maladaptation” has been used to describe both the state and the process. Maladaptation has many potential causes, including genetic constraints, genetic drift, and mutational load (Brady et al. 2019a, b). Here, we focus on the somewhat counterintuitive possibility that selection itself may drive the process of maladaptation (Wright 1942; Lande 1976; Svensson and Connallon 2019).

To understand how selection may lead to maladaptation, it is instructive to explore Fisher’s (1930) fundamental theorem of natural selection, which has often been the focus of theoretical and empirical treatments of the evolution of population fitness. In its most basic form, the fundamental theorem states that the partial change in population mean fitness due to natural selection is equal to a population’s additive genetic variance for fitness (Price 1972a; Frank and Slatkin 1992; Burt 1995; Frank 1997; Hendry et al. 2018; Bonnet et al. 2019). Interpreted on its own, the partial change described by the fundamental theorem would appear to suggest that natural selection cannot lead to maladaptation, as the additive genetic variance in fitness can never be less than zero. Later exegeses of the fundamental theorem showed that Fisher’s partial change means something very specific, namely, the change in population fitness caused by allele frequency changes, while holding constant any changes in the average effects of alleles (Price 1972a; Ewens 1989; Okasha 2008; Ewens and Lessard 2015). Any shifts in population fitness driven by change in the average effects of alleles, or by other sources such as environmental change, frequency dependence, dominance, gene interactions, or other nonadditive effects are relegated to a component referred to as the “deterioration of the environment” (Price 1972a; Frank and Slatkin 1992; Hadfield et al. 2011). Fisher (1930) viewed environmental deterioration as typically acting in opposition to natural selection, reducing mean fitness each generation and facilitating continuous phenotypic evolution. Using the metaphor of the adaptive landscape, environmental deterioration should cause peaks to move slightly each generation, thwarting a population’s ascent to optimal mean fitness. In some cases, however, the deterioration of the environment may be extensive enough to overwhelm adaptation, leading to an overall decrease in mean fitness each generation and resulting in maladaptation (Brady et al. 2019a). Because selection can underlie environmental deterioration, e.g. through frequency dependence or socially mediated effects, selection may thus be the root cause of some maladaptation (Wright 1942, 1969; Lande 1979; Brady et al. 2019a; Fisher and McAdam 2019; Svensson and Connallon 2019).

Social interactions among conspecifics may be particularly likely to lead to reductions in population fitness and thus maladaptation (Matsuda and Abrams 1994; Kokko and Brooks 2003; Hadfield et al. 2011; Brady et al. 2019b; Fisher and McAdam 2019; Svensson and Connallon 2019; Henriques and Osmond 2020). While social interactions may often enhance adaptation when the interests of social partners are aligned (Henriques and Osmond 2020), they may lead to maladaptation when competitive traits that increase individual fitness at the expense of others evolve (Wright 1969; Matsuda and Abrams 1994; Webb 2003). In extreme cases, selection involving interspecific competition may decrease population mean fitness enough to lead to population extinction, a phenomenon known as “self-extinction” (Matsuda and Abrams 1994), “evolutionary suicide” (Gyllenberg and Parvinen 2001), or “Darwinian extinction” (Webb 2003).

Several recent models have explicitly considered social interactions in the context of Fisher’s fundamental theorem (Bijma 2010b; Hadfield et al. 2011; Queller 2014; Fisher and McAdam 2019). Bijma (2010b) developed a model that included heritable influences of an individual’s social environment, or indirect genetic effects (IGEs), on fitness. Bijma’s approach showed that the partial increase in population fitness predicted by Fisher’s fundamental theorem is equivalent to additive genetic variance in inclusive fitness, supporting earlier conclusions that selection tends to lead to the optimization of inclusive fitness (Hamilton 1964; Grafen 2006). Bijma identified that the deviation between change in inclusive fitness and change in population mean fitness may stem from the deterioration of the environment caused by social competition. Hadfield et al. (2011) further showed that earlier models of decreases in population fitness due to intraspecific competition (Cooke et al. 1990) produced an effect equivalent to Fisher’s (1930) deterioration of the environment. Queller (2014) elaborated on these results, explicitly modeling the decrease in fitness caused by antagonistic social interactions and discussing conditions in which the change in mean population fitness would diverge from the change in inclusive fitness. Finally, Fisher and McAdam (2019) showed that such socially mediated decreases in population fitness may be viewed as arising from a negative covariance between direct and indirect genetic effects on fitness. Although the direct component of fitness should always increase (assuming that the average effects of alleles do not change), there can be correlated changes in how individuals influence the fitness of others, perhaps due to competition for limited resources. An increase in the detrimental effects on the fitness of others due to a negative direct-indirect genetic covariance is equivalent to a deterioration of the social environment and can potentially be large enough to overwhelm the adaptive effects of natural selection, leading to maladaptation.

Despite this earlier work, we lack a clear rule identifying when socially mediated selection should lead to maladaptation. Here, we develop such a rule by reformulating earlier results in terms of social selection, which is defined as selection mediated by phenotypes in the social environment (West-Eberhard 1979; Wolf et al. 1999). Social selection may be distinguished from nonsocial selection via regression analysis, where the nonsocial selection gradient is the partial regression of an individual’s fitness on its own traits and the social selection gradient is the partial regression of an individual’s fitness on traits of its social partners. Formulating a model of adaptation in terms of these gradients allows us to make explicit connections between phenotypic evolution and the predicted change in population fitness due to selection. Our approach also generates expressions written in terms of parameters that can be estimated in natural populations, providing simple empirical tests for socially mediated maladaptation that can be employed in natural populations.

## Social selection and the evolution of fitness

To determine when social interactions should lead to the evolution of maladaptation, we model the predicted change in fitness in a single generation in response to two types of phenotypic selection. Nonsocial selection (**β**_N_) measures the effect of a focal individual’s own traits on its own fitness, while social selection (**β**_S_) measures the effect of the traits of social partners on a focal individual’s fitness. For the purposes of the model, we define maladaptation as an evolved decrease in fitness (*w*) from one generation to the next. We make no assumptions about fitness other than it can be decomposed into a heritable component (or breeding value, *A*_*w*_) and a residual component (*e*_*w*_), which are assumed to be uncorrelated. This is equivalent to considering a partial change in fitness that ignores dominance, epistasis, and changes in the mean effect of alleles, following Fisher (1930). Our model differs, however, in incorporating social effects into the breeding value as in models of IGEs (Moore et al. 1997; Bijma and Wade 2008; McGlothlin et al. 2010; Fisher and McAdam 2019). This allows us to isolate the effects of social selection on the partial change in mean fitness. The absolute fitness of an individual is often operationally defined in quantitative genetics as its lifetime reproductive success, with the generational dividing line placed at the zygote stage (Arnold and Wade 1984; Wolf and Wade 2001). Because our model involves only a single generation of evolutionary change, lifetime reproductive success is a suitable surrogate for absolute fitness in the derivation that follows. Extension of our results to long-term predictions would require incorporating an explicit model of population regulation (Hendry 2017), which is beyond the scope of this paper.

For simplicity, our model tracks relative fitness (*w*), which we define as absolute fitness divided by the population mean fitness. Relative fitness can also be written as a sum of a heritable component and a residual component, or

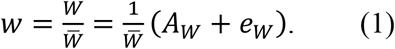

The heritable component of relative fitness, or its total breeding value, is just the first term on the right-hand side of Equation 1, or

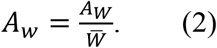

In what follows, we assume that before selection, the mean breeding value for relative fitness equals population mean relative fitness 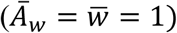, while the residual component has a mean of zero. To extend our model beyond a single generation, additional assumptions about the nature of this breeding value are necessary. For example, the breeding value for relative fitness could be defined with respect to fitness at some arbitrary time point, with the phenotypic value of relative fitness rescaled each generation as mean fitness evolves.

Equation 2 indicates that for a single generation, change in the heritable component of absolute fitness 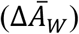 will be proportional to the change in the heritable component of relative fitness multiplied by mean absolute fitness 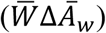. To simplify notation, we focus on a single generation of change only and write the partial change in relative fitness due to heritable changes simply as 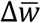. From the Price (1970, 1972b) equation, this component of change can be written as the covariance of the total breeding value for relative fitness with relative fitness itself, or

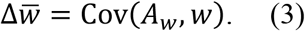

We note that Equation 3 does not include changes in mean fitness that are due to environmental or non-additive genetic effects, and in the absence of social effects, is equal to the additive genetic variance in relative fitness, or equivalently, the evolvability of fitness (Fisher 1930; Price 1972a; Frank and Slatkin 1992; Houle 1992; Burt 1995; Orr 2009; Hendry et al. 2018). In what follows, we operationally define a generation of adaptive change as a positive value of 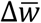 and a generation of maladaptive change as a negative value of 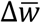.

When there are social effects on fitness, the covariance in Equation 3 will not equal the additive genetic variance in relative fitness and thus may take on negative values. Previous models have decomposed the total breeding value for fitness into direct and indirect (social) components (Bijma 2010b; Fisher and McAdam 2019). Here we link this decomposition to phenotypic selection, which ultimately underlies genetic variance for fitness and, in most cases, is easier to estimate in nature. Our goal is to describe the conditions in which a combination of nonsocial and social selection should lead to maladaptation, or an evolved decrease in mean fitness 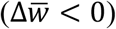, in a single generation. To this end, we first express relative fitness as a function of focal and social phenotypes,

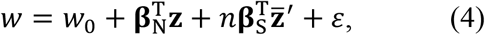

(Wolf et al. 1999; McGlothlin et al. 2010) where the intercept (*w*_0_) is a population parameter and the error term (*ɛ*) has a mean of zero. In Equation 4, **Z** is a vector of a focal individual’s traits, and 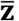′ is a vector containing the mean phenotypes of that individual’s social interactants. The superscript T designates matrix transposition, which here indicates that a column vector is to be written as a row vector. The product of each selection gradient and phenotype vector is thus a scalar, each of which indicates the total fitness effects of an individual’s own traits and those of its social partners, respectively.

Mean relative fitness can calculated by taking the expectation of Equation 4,

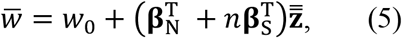

where 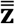 is the vector of population mean phenotypes. Equation 5 assumes that all individuals have *n* social partners, but if social group size varies and *n* and 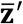 are uncorrelated *n* can be rewritten as 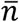. We then define an individual’s total breeding value for relative fitness (*A_w_*) as

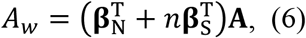

where **A** represents an individual’s vector of total breeding values for the phenotypes represented by **z**. This definition of *A*_*w*_ decomposes an individual’s effect on population mean fitness into effects of individual traits. Through the two selection gradients, trait effects are further decomposed into nonsocial and social components. Therefore, these components (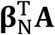 and 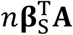, respectively) are the trait-based equivalents of the total direct and indirect genetic effects on fitness (Fisher and McAdam 2019).

Because breeding values for fitness are now a function of phenotypic breeding values, the change in relative fitness due to one generation of phenotypic selection can be found by substituting Equation 6 into Equation 3:

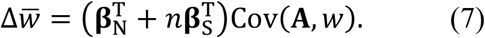

From the Price (1970, 1972b) equation, the covariance between the phenotypic breeding value and fitness equals the total phenotypic change due to selection 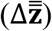, so Equation 7 can be rewritten as

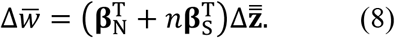

This phenotypic change can itself be expanded as a function of selection, direct and indirect genetic effects, and relatedness (Bijma and Wade 2008; McGlothlin et al. 2010). When social selection, relatedness, and IGEs are absent, our equation is equivalent to the result derived by Lande (1979) showed that when social effects are absent and when selection is frequency-independent, 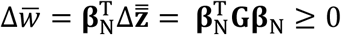, where **G** is the additive genetic (co)variance matrix. Therefore, without social effects or frequency dependence, selection is not expected to lead to a decrease in fitness.

By a simple rearrangement of Equation 8, we can obtain an informative result. As written, the right-hand side of Equation 8 is the dot product of two vectors, the first of which is a simple sum of nonsocial and social selection gradients (**β**_N_ + *n***β**_S_) and thus represents the total phenotypic effect of traits on fitness in a population, and the second of which represents evolutionary response to selection 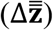, or how these phenotypic effects translate into genetic change. The dot product of two vectors is proportional to the cosine of the angle between them (*θ*), so we can rewrite the right-hand side of Equation 8 as

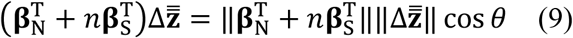

where the double vertical bar notation indicates the vector norm (i.e., the magnitude of the vector, which is always a non-negative scalar). This equation allows us to view the potential for maladaptation geometrically, as a misalignment between selection and the evolutionary response to selection. The cosine function is bound by −1 and 1, so we consider the space where *θ* is between 0° (cos *θ* = 1; vectors completely aligned) and 180° (cos *θ* = −1; vectors completely misaligned). It then follows that selection will lead to adaptation 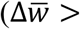 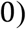 when 0° ≤ *θ* < 90° and maladaptation 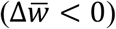 when *θ* > 90°, with the maximum decrease in fitness occurring when *θ*= 180° (cf. Hadfield and Thomson 2017). In other words, an alignment between the phenotypic selection and phenotypic evolutionary change leads to adaptation, while misalignment leads to maladaptation.

Lande’s (1979) result that 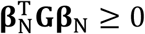 indicates that in the absence of social effects, natural selection and evolutionary change are always aligned (or at least orthogonal) and that the partial change in fitness due to selection should never be negative. However, Equation 9 suggests that when social effects exist, selection and evolutionary change may be misaligned (Fisher and Pruitt 2019), which predicts a decrease in population fitness. Fisher and McAdam (2019) discuss a number of scenarios involving social interactions that can lead to ongoing trait evolution with zero change in population fitness or even maladaptation. For example, population may be limited by the availability of refuges, food, mates, or some other resource, leading to social competition. When competition occurs, the same traits that lead to individual benefits will reduce the success of competitors, which should tend to create nonsocial and social selection gradients in opposing directions. This effect has been identified as a source of evolutionary constraint and has been referred to as both the “intraspecific Red Queen” and the “treadmill of competition” (Rice and Holland 1997; Wolf 2003; Wilson et al. 2009, 2011; Wilson 2014).

To understand the conditions that may lead to a misalignment between selection and evolutionary response, it is instructive to expand Equation 8 to include the relationship between genes and phenotype. In doing so, we examine the simplest case, the evolution of a single trait with social interactions occurring between pairs of individuals (i.e.,*n* = 1). In this case, the condition for maladaptation simplifies to

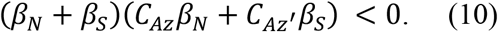

where the two selection gradients, *β*_*N*_ and *β*_*S*_, are now expressed as scalars, *C*_*Az*_ is the covariance between an individual’s breeding value and its own phenotype, and *C*_*Az′*_ is the covariance between an individual’s breeding value and its partner’s phenotype (McGlothlin et al. 2010). These covariance terms in turn depend on genetic variance (*G*), relatedness (*r*), which we define as the correlation between a focal individual’s additive genetic value and that of its social partner, and the interaction coefficient *ψ*, which measures the effect of a social partner’s trait on phenotypic expression in the focal individual. When traits are heritable, *ψ* represents the strength and direction of trait-based IGEs (Moore et al. 1997). Expanding, the two covariances in Equation 10 can be written as

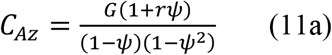

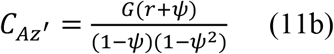

(McGlothlin et al. 2010). The denominators in Equation 11 represent IGE feedback effects that occur because the effects on trait expression are bidirectional (Moore et al. 1997). Substituting these terms into Equation 10 and rearranging, we can express the conditions for maladaptation as

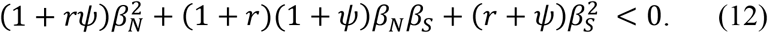

Because *r* and *ψ* are normally bound by −1 and 1, the occurrence of maladaptation depends on two key quantities, *β_N_β_S_* and *r* + *ψ*. These two quantities respectively represent conflict in the levels of selection and the two pathways, relatedness and IGEs, by which interacting phenotypes may become correlated. On their own, or in concert, these two quantities can be sufficiently negative to overwhelm the first term of on the left-hand side of Equation 12, which is always positive. Stated another way, maladaptation is most likely to occur when nonsocial and social selection are in conflict and/or when the phenotypes of interacting individuals are negatively correlated due to the combination of relatedness and IGEs.

In order to simplify our formulation and identify general situations when social selection can lead to maladaptation, we can remove IGEs by setting *ψ* = 0. Equation 12 therefore becomes

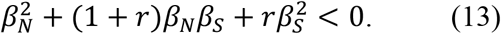

Figure 1 shows when the condition in Equation 13 is satisfied for two different scenarios: when a trait confers an individual benefit (*β*_*N*_ > 0) and when a trait is costly to self (*β*_*N*_ < 0). Figure 1 displays only the regions of parameter space where an evolutionary increase in the trait occurs 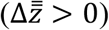. Cases where stasis or a trait decrease is predicted are interpreted as a trait being unable to increase by natural selection and are shown as empty regions in Figure 1. Figure 1 shows that maladaptation can occur in two different situations. The most likely situation occurs when a trait may either be beneficial to self and harmful to others (*β*_*N*_ > 0, *β*_*S*_ < 0), which represents selfishness or competition (Figure 1A, left side). This scenario corresponds to classic results showing that intraspecific competition may lead to selection that decreases population fitness (Cooke et al. 1990; Matsuda and Abrams 1994; Gyllenberg and Parvinen 2001; Webb 2003; Fisher and McAdam 2019; Svensson and Connallon 2019; Henriques and Osmond 2020). As a selfish or competitive trait increases by natural selection, the trait imposes fitness costs on an individual’s social interactants. If the magnitude of *β*_*S*_ is large enough, this will lead to an evolutionary decrease in population mean fitness (Figure 1A). In Equation 13, this effect is caused primarily by the second term. When *β*_*N*_*β*_*S*_ < 0 and is relatively large in magnitude, social costs will tend to overwhelm individual benefits and fitness will decrease even as the trait itself evolves. Relatedness interacts with this effect in a complex way. When *r* > 0, relatives tend to pay the social cost. As relatedness increases, eventually the accumulation of costs to relatives means that the trait cannot increase by natural selection (Figure 1A, upper left corner). When *r* < 0, an individual is more likely to interact with nonrelatives than would be expected by chance. As *r* becomes more negative, the third term in Equation 13 becomes important, intensifying the effect of maladaptation (Figure 1A, bottom left corner).

**Figure 1.**
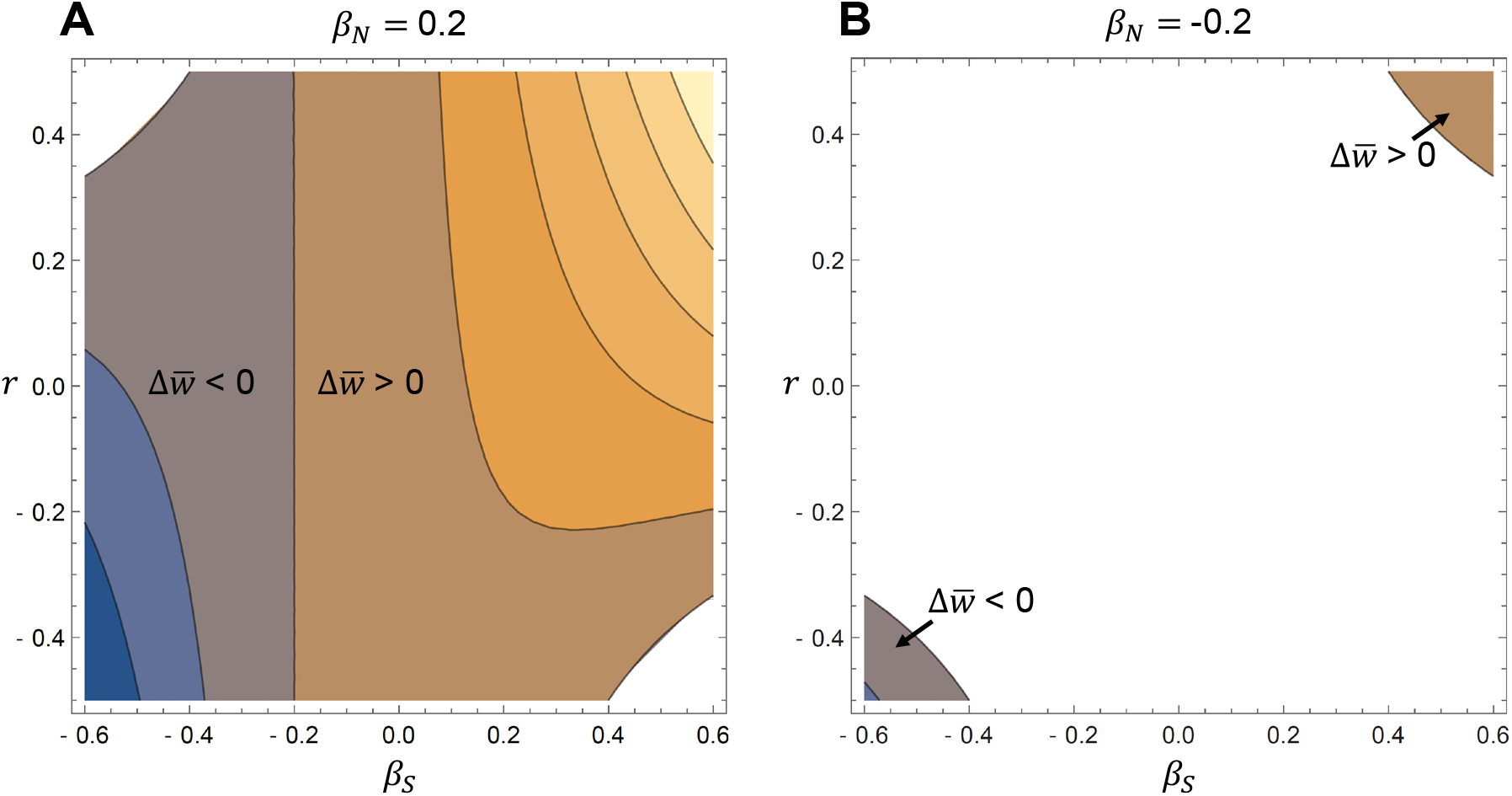
Conditions for the evolution of maladaptation derived from Equation 13. Here we consider a single trait under positive (A) or negative (B) nonsocial selection (*β*_*N*_). Regions where selection leads to an increase in fitness are shown in orange, and regions where fitness decreases are shown in blue. Only regions where the trait can increase via natural selection are shown. Contour lines represent increments of 0.066. Maladaptation occurs when social selection (*β*_*S*_) is negative and outweighs individual benefits (A, left) or in the evolution of spite (B, bottom left corner).

When *β*_*N*_ < 0, the regions in which the trait may evolve are limited (Figure 1B). Hamilton’s (1964) rule predicts the evolution of a costly trait in two scenarios. The classic case is altruism, where a costly trait leads to a social benefit (*β*_*S*_ > 0). This can only occur when relatedness is high enough to give a product with social benefits that outweighs individual costs (Figure 1B, upper right corner). The evolution of altruism leads to an increase in population fitness as would be expected from previous work (e.g. Bijma 2010b; but see Henriques and Osmond 2020 for one scenario where the evolution of altruism may lead to a decrease in fitness). An interesting case arises when *β*_*N*_ and *β*_*S*_ are both negative, which corresponds to the selection expected for a spiteful behavior. If *r* is negative and of large enough magnitude, a spiteful trait can evolve (Gardner and West 2004), which in our model leads to an evolved decrease in fitness as both individuals and social partners suffer a fitness cost (Figure 1B, bottom left). From Equation 13, it is evident that this effect arises when the third term is sufficiently negative to overwhelm the other two terms, which will be positive in this scenario.

Although Equation 13 is written for pairwise interactions, it can be generalized to larger social groups by replacing *β*_*S*_ with *nβ*_*S*_, where *n* is the number of groupmates excluding the focal individual. When doing so, the condition for the evolution of maladaptation in Equation 13 is equivalent to results derived from variance-component based models that include IGEs for fitness (Bijma 2010b; Fisher and McAdam 2019). In such models, the trait under selection is fitness itself, so 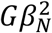 would represenhte dt irect genetic variance for fitness, 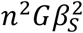 would represent indirect genetic variance for fitness, and *Gβ_N_β_S_* would represent the direct-indirect covariance. Fisher and McAdam (2019) noted that the most likely scenario for maladaptation occurs when the direct-indirect covariance is negative and further from zero than the direct genetic variance for fitness. Our trait-based model corresponds to this conclusion and further shows that this scenario is most likely when social selection is strong and acting in opposition to nonsocial selection.

To further simplify, we can consider the special case with no relatedness, in which Equation 13 reduces to:

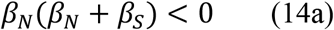

or

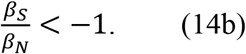

Thus, when non-relatives interact in the absence of IGEs, maladaptation is driven only by the magnitude and sign of the selection gradients. As discussed above, the most likely scenario satisfying the condition in Equation 14 is strong social competition, where positive values of a trait are beneficial to self (*β*_*N*_ > 0) but harmful to others (*β*_*S*_ < 0), and the magnitude of the harm outweighs the benefit (|*β*_*S*_| > |*β*_*N*_|). As the selfish trait increases each generation, the population fitness declines. If the trait were harmful to self (*β*_*N*_ < 0) and beneficial to others (*β*_*S*_ > 0) however, which represents an altruistic trait, the trait would be removed by natural selection. This is because in the absence of relatedness and IGEs, social selection would not contribute to a response to selection and altruism cannot evolve (Bijma and Wade 2008; McGlothlin et al. 2010).

## Empirical applications

The results of our model support previous work indicating that selection arising from social interactions can decrease population mean fitness, particularly in the case of competition (Wright 1969; Matsuda and Abrams 1994; Gyllenberg and Parvinen 2001; Webb 2003; Hadfield et al. 2011; Fisher and McAdam 2019; Svensson and Connallon 2019; Henriques and Osmond 2020). However, because our results are given in terms of quantitative genetic selection gradients, the parameters of our model can be readily estimated in natural populations, leading to empirical tests of the contribution of phenotypic selection to maladaptation. Nonsocial and social selection gradients may be estimated in natural populations via a modification of the standard Lande-Arnold (1983) method by incorporating social phenotypes into a multiple regression as in Equation 5 (Wolf et al. 1999). Once selection gradients have been estimated, they may be compared to the vector of phenotypic responses to selection (either predicted or observed) using Equation 8 or 9 to test whether selection is contributing to maladaptation in a given generation. Below we outline how different types of data may be used for such tests.

The most straightforward empirical test of the role of social selection in maladaptation would involve studying a population for two generations. In the first generation, measurements of the phenotypes of focal and social individuals and relative fitness from the same individuals (preferably measured as relative lifetime reproductive success) would be used to estimate nonsocial and social selection gradients. In the second generation, measurements of the same phenotypes as in the previous generation would be used to estimate evolutionary change 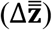. Then, Equation 8 or 9 could be used to test for maladaptation by taking the product or the angle between the vector of total selection and the vector of evolutionary change.

If quantitative genetic data are available, the predicted response to selection may be substituted for observed evolutionary change. For phenotypes involving social interactions, the total predicted evolutionary change involves estimates of the additive genetic (co)variance matrix **G**, IGEs, and relatedness (Bijma and Wade 2008; McGlothlin et al. 2010). IGEs may be measured using a variance-component model or a trait-based model, which make equivalent predictions of evolutionary response to selection (McGlothlin et al. 2010). However, because IGEs are difficult to measure in practice (McGlothlin and Brodie 2009; Bijma 2010a), this approach may be infeasible in many cases if IGEs cannot be ignored. Alternatively, the genetic covariance between traits and fitness, if estimable from the data, can be used to predict phenotypic evolutionary change (Morrissey et al. 2010; Morrissey et al. 2012). Using predicted evolutionary change is beneficial for two reasons. First, it helps disentangle genetic shifts in the population mean from environmental change, which might bias the method outlined above. Second, it may allow the application of Equation 8 or 9 when phenotypic data are not available from the offspring generation.

In the absence of quantitative genetic data or observed evolutionary change, it is possible to use a phenotypic version of Equation 8 or 9 as a proxy to investigate whether social interactions can lead to maladaptation. In place of 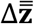, one may substitute the realized selection differential **s** = **Pβ_N_** + *n***C^I^β_S_**, where **P** is the phenotypic (co)variance matrix and **C^I^** is the covariance matrix of the phenotypes of interacting individuals (Wolf et al. 1999). This vector represents the total effect of phenotypic selection on the mean phenotype of the population before accounting for genetic transmission. Both **P** and **C^I^** include effects of relatedness and IGEs (Moore et al. 1997; Wolf et al. 1999), allowing them to act as surrogates for their genetic counterparts. Relatedness and IGEs are particularly important for **C^I^**: random interactions with no IGEs should lead to elements of **C^I^** that are close to zero, while both relatedness and IGEs may lead to either positive or negative values (Wolf et al. 1999). The disadvantage of this approximation, however, is that environmental sources of (co)variance also contribute to both **P** and **C^I^**, making **s** potentially deviate from 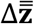. If such deviations are primarily in magnitude, the purely phenotypic version of Equation 9 will still be a useful to indicate whether social interactions will alter the change in mean fitness, but if environmental effects lead **s** and 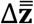 to differ in direction, Equation 9 may be misleading. We argue that the benefits of such phenotypic approximations are likely to outweigh their costs. Using a phenotypic version of Equation 9 will allow a much wider application of our results to systems where collecting quantitative genetic data is difficult or impossible, providing a much broader picture of the potential importance of social selection for maladaptation at the expense of a degree of accuracy.

Finally, we note that the simplest test of the contribution of social selection to maladaptation can be conducted with selection gradients alone. Equation 14 shows that when relatedness and IGEs are absent, the contribution of social selection on a single trait to maladaptation depends only on the signs and relative magnitude of the two selection gradients. Specifically, a single trait can be considered to contribute to maladaptation if social selection is stronger than natural selection and opposite in sign. Further, as shown in Figure 1, the selection gradients dominate the direction of the predicted change in population fitness unless the covariance between interactants is strong. Thus, selection gradients alone may be used as a rough estimate of whether a single trait contributes to maladaptation. Data from competition. Fisher and Pruitt (2019) reviewed studies in which nonsocial and social selection were both measured. These two selection gradients often differed in sign, indicating conflict, and in a few cases conflicting social selection was stronger than nonsocial selection. Intriguingly, one such case involved selection for arrival date in a territorial songbird, hinting that social competition for nest sites may be individually beneficial but detrimental to population mean fitness (Farine and Sheldon 2015).

## Inclusive fitness

Our results align with previous studies showing that natural selection does not always lead to an increase in population mean fitness when social interactions occur. While we have equated a decrease in population mean fitness with maladaptation, this view is not universally held. For example, Gardner and Grafen have argued that because selection tends to optimize inclusive fitness, adaptation should be viewed from the perspective of inclusive fitness rather than population mean fitness (Grafen 2006; Gardner 2009; Gardner and Grafen 2009). As defined by Hamilton (1964), inclusive fitness removes fitness effects of social partners but adds in fitness effects on social partners, weighted by relatedness (or when IGEs are present, by a function of relatedness and IGEs, McGlothlin et al. 2010). Thus, although social context is important, inclusive fitness effects are defined as arising from the focal individual. Inclusive fitness theory predicts that individuals are expected to act as if they are maximizing inclusive fitness, which provides the rationale for using it to define adaptation (Hamilton 1964; Dawkins 1982; Grafen 2006; Gardner and Grafen 2009; Bijma 2010b). Bijma’s (2010b) results, which show that the partial increase in population mean fitness predicted by Fisher’s fundamental theorem is identical to the predicted increase in inclusive fitness, support this perspective.

By defining inclusive fitness as a function of nonsocial and social selection as in (McGlothlin et al. 2010) and following the logic of Equations 1–4, the change in mean inclusive relative fitness is predicted by

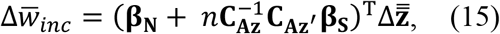

which, although expressed in different notation, is equivalent to Equation 37 of Hadfield and Thomson (2017). In Equation 15, the covariance matrices are the multivariate equivalents of the covariances in Equation 7 (McGlothlin et al. 2010). Hadfield and Thomson (2017) show that, except in the case of singularity of **G**, Equation 15 is always positive, indicating that relative inclusive fitness always increases even when population mean fitness does not. Comparing Equation 15 to Equation 8 shows why inclusive fitness should always increase, even while population mean fitness may decrease. As shown in Equation 8, the contribution of social selection to the change in population mean fitness does not depend on the matrix **C_AZ′_**. Because this matrix determines the contribution of social selection to phenotypic evolutionary change, the total effect of selection on fitness can diverge from evolutionary response, causing an evolutionary decrease in population mean fitness. However, in Equation 15, **C_AZ′_** determines the effect of social selection on both change in inclusive fitness and evolutionary response to selection, meaning the two should always be aligned. Organisms should therefore act to maximize their inclusive fitness, even if this means reducing the fitness of others and ultimately lowering population mean fitness.

## Conclusion

Here we show that previous treatments of maladaptation via selection (e.g. Fisher and McAdam 2019; Svensson and Connallon 2019) may be usefully viewed from the perspective of social selection. When social selection is strong and in conflict with nonsocial selection, and/or when there is strong negative covariance in the phenotypes of interacting individuals, the partial increase in population mean fitness predicted by Fisher’s fundamental theorem may be overwhelmed by a deterioration of the social environment, leading to maladaptation. Interestingly, the conditions for maladaptation are identical to the conditions previously identified for responses to selection that occur in an opposing direction to selection itself (Fisher and Pruitt 2019). Indeed, the crucial test for whether a group of traits contributes to maladaptation involves the misalignment of selection and its predicted response.

Our results also provide further rationale for measuring social selection in the wild (Wolf et al. 1999). Although estimates of social selection are accumulating (Formica et al. 2011; Farine and Sheldon 2015; Fisher and Pruitt 2019), they are still rare. Most arguments advocating for the measurement of social selection consider it in the context of altruism and Hamilton’s rule (McGlothlin et al. 2014), but the fact that the strength of social selection may reflect whether a population is undergoing adaptation or maladaptation indicates that it may be equally important in the context of social competition.

## Acknowledgments

We thank Kim Hughes and Anjanette Baker for organizing the symposium and editing this theme issue. We thank Andrew Hendry, Andrew McAdam, Josef Uyeda, and two anonymous reviewers for helpful comments on the manuscript.

